# Novel genetic model of pediatric Diffuse Intrinsic Pontine Glioma in *Drosophila melanogaster*

**DOI:** 10.1101/2023.07.10.548387

**Authors:** Carmen de Pablo, Sergio Casas-Tintó

**Affiliations:** IIER-Instituto de Salud CarlosIII, Majadahonda, Spain

**Keywords:** midline glioma, H3K27, MYC, brain tumor, cancer, DIPG, *Drosophila*, Diffuse Midline Glioma, Glioblastoma, H3K27M, glia

## Abstract

Diffuse Intrinsic Pontine Glioma (DIPG) is a lethal pediatric type of brain tumor that grows in the bm and originated from glial cells. Its location and infiltrative nature impede surgical resection and make the treatment difficult and low effective. In consequence, affected children have a short life expectancy of 12 months. The most frequent mutation is a substitution of lysine to methionine at residue 27 of histone H3 (H3K27M). Secondary mutations in additional genes, including *Myc*, are required for the malignancy of glial cells. The lack of studies and tumor aggressiveness make it necessary to generate new experimental models that reproduce the fundamental aspects of the disease and allow to expand the knowledge about DIPG.

*Drosophila melanogaster* presents advantages as an experimental model and stands out for its genetic tools, easy handling, and great genetic and cellular homology with humans. *Drosophila* has contributed to the investigation of different diseases, including glioblastoma (GB) and neurodegenerative diseases as Alzheimeŕs or Parkinsońs. Here we present a new genetic model of DIPG generated in *Drosophila melanogaster*. It is based on the overexpression of *H3K27* and *Myc* in glial cells that produce an increase in the number of glial cells in the ventral nerve cord and the expansion of glial membranes in early developmental stages. However, this novel DIPG model does not produce tumoral features in adult brains, in line with the pediatric nature of this disease. We have evaluated the activation of different signaling pathways active in other glial tumors, in this model of DIPG. The results show that, unlike GB, JNK pathway is not upregulated in DIPG, and it is not determinant for the progression of DIPG. Besides, glial cells in the DIPG model accumulate MMP1 and MMP2 and increase the accumulation of Liprin-γ, previously associated to the formation of synaptic structures in GB cells. The results show that DIPG is a unique entity that differs from other high-grade gliomas such as GB and will require of a different therapeutic approach.

## INTRODUCTION

Tumors or neoplasms are abnormal masses of tissue that grow uncontrollably, excessively, autonomously, and irreversibly. According to its clinical behavior we find benign tumors, which do not generate secondary growth, and malignant tumors, of an infiltrative nature and with the capacity to generate secondary implants or metastasis.

Tumors of the Central Nervous System (CNS) represent the seventh neoplasia in frequency within adult population (Louis et al. 2021). They are those that develop in the brain, spinal cord or meninges. Based on the histological characteristics and molecular parameters, the WHO differentiates four grades of CNS tumors according to their aggressiveness, grade I is the less aggressive and IV the most aggressive.

Gliomas are the most common tumors of the CNS, originating from neoplastic glial cells. According to the WHO classification we find 4 types of gliomas: diffuse adult-type gliomas, pediatric-type diffuse low-grade gliomas, pediatric-type diffuse high-grade gliomas, and circumscribed astrocytic gliomas (Louis et al. 2021). In this classification the authors indicate the clinical and molecular differences between diffuse gliomas present mainly in adults (“adult type”) and those present mainly in children (“pediatric type”), diffuse low-grade gliomas of pediatric type and diffuse high-grade gliomas of the pediatric type.

### High Grade Supratentorial Gliomas

High-grade gliomas (HGG, *Supratentorial High-grade Gliomas)* represent a spectrum of diseases, with histopathological features shared between childhood and adult tumors (Louis et al. 2021), as well as mutations in the same canonical cancer pathways such as the receptor tyrosine kinase (RTK)/RAS/PI3K, the TP53 pathway, and the RB pathway, although the effectors most affected by the mutation vary, between childhood and adult tumors. Even though the effectors commonly affected by mutations differ between pediatric and adult tumors (Sturm et al. 2014).

Recent molecular profiling data demonstrate that childhood tumors are biologically distinct from adults and suggest that there are unique molecular processes that drive tumorigenesis in the developing brain, which vary by age and anatomical location. These distinctions reflect the need for very different therapeutic approaches to effectively counter underlying genetic mutations (Jones & Baker 2014).

### Diffuse midline glioma

Diffuse intrinsic pontine glioma (DIPG) or diffuse midline glioma has been classified as a grade IV entity within the Pediatric-type HGG (Louis et al. 2021). It is a pediatric brain stem glioma that originates from the ventral pons, accounts for 75-80% of brainstem tumors in children, it has a peak incidence in middle childhood and a life expectancy of less than 1 year (Lapin et al. 2017; Wu et al. 2014).

Diagnosis is based on the diffuse infiltration of the midline crossing, typical histological features, and the presence of alterations in residue 27 of Histone H3 (H3K27) (Louis et al. 2016, 2021).

DIPG cannot be removed surgically due to its location and the infiltrative nature of the disease. Palliative radiotherapy is the only therapy, although it only provides temporary improvements on neurological and radiological function (Lapin et al. 2017; Wu et al. 2014). And despite the trials, DIPG are still considered fatal pediatric tumors without cure (Lapin et al. 2017).

### Genetic basis

The acquisition of samples of primary tumors has facilitated the elaboration of genomic profiles and advances in the knowledge of key oncogenic factors. This has favored the identification of the genes responsible for the appearance and progression of this type of tumor.

Proteomic and transcriptomic analysis of DIPG suggests that it is a unique type of glioma that shares biological similarities with HGGs, such as glioblastoma multiforme (GB) (Jones & Baker 2014).

Likewise, there is a great intra- and intertumoral heterogeneity within the markers that define the disease, making it difficult to develop effective therapeutic strategies (Hoffman et al. 2016). Analysis by sequencing of DIPG samples showed mutations, including single nucleotide variations, insertions and deletions, and structural variations.

### Histone H3

The most relevant genetic characteristic of this tumor type is the presence of mutations in *histone H3*, present in 80% of DIPG cases (Lapin et al. 2017).

Histone mutations in DIPG are subclassified into histone H3.1 (H3.1) or histone H3.3 variants. (H3.3), encoded by the genes *HIST1H3B* and *H3F3A,* respectively. There are clinical features specific for each variant. Histone H3.1 is associated with a median overall survival of 15 months and a lower presence of metastases, while histone H3.3 mutations are related to a median overall survival of 9 months and less response to radiotherapy. In both cases, DIPGs with mutations in *H3* are associated with poorer outcomes compared with tumors with wild type *histone H3* (Lapin et al. 2017).

The specific mutation in histone H3 is caused by the substitution of a lysine to methionine at the residue 27 (H3K27M). H3K27 is a key center for transcriptional regulation: acetylation (H3K27ac) is associated with active enhancers, whereas trimethylation (H3K27me3) represses transcription. This methylation is carried out by the Polycomb repressive complex 2 (PRC2) (Nagaraja et al. 2019). H3K27M has a greater affinity for PRC2, inhibiting it and therefore reducing H3K27me3. However, H3K27M also causes an increase in H3K27me3 in specific tissues, suggesting that both expression and repression of abnormal genes is important (Pajovic et al. 2020).

The tumorigenesis-initiating role of oncohistone in DIPG relies on remodeling of the transcriptome: H3K27M induces epigenetic changes that activate multiple members of the RAS/MAPK and MYC cascade, as well as their downstream transcriptional targets. However, the activation during early development of the epigenetic pathway of H3K27M is not sufficient to produce tumorigenesis in DIPG, and it is later reinforced by pathway-activating gene mutations (Pajovic et al. 2020). This evidence explains the coexistence of these mutations in human DIPG.

### Myc

*C-myc* is one of the most amplified oncogenes in human cancer and plays a fundamental role in tumor transformation. This gene encodes the protein MYC, a transcription factor that regulates cell proliferation, differentiation, apoptosis, and cell migration. Rising levels of MYC drive tumor initiation, progression, and recurrence, and are necessary for the tumor maintenance (Hutter et al. 2017).

One of the transcriptional features of H3K27M tumors is the epigenetic activation of MYC. Loss of H3k27me3 induces overexpression of MYC target genes in DIPG (Pajovic et al. 2020). MYC amplifications and overexpression occur in 20% of the cases. MYC alterations are present in DIPG and act as genome-wide transcriptional regulators (Lapin et al. 2017).

### Glioblastoma

The most common type of glioma is glioblastoma multiforme (GB), classified by the WHO as a astrocytic and diffuse oligodendroglial grade IV brain tumor. It is the most aggressive and lethal brain tumor, with an incidence of three per 100,000/year. The median survival of patients with GB is 12-15 months, with less than 5% chance of survival after 5 years (Gallego 2015; Louis et al. 2016; McGuire 2016; Rogers et al. 2017). Temozolomide (TMZ) is the only treatment for GB, however, recent studies restrict its use in patients with GB based on the methylation status of *methylguanine DNA methyltransferase (MGMT)* (Wick et al. 2018). Besides, the Genetic and molecular heterogeneity complicates the diagnosis and treatment of these tumors.

Recent studies have revealed that GB cells extend membrane ultra-long tubes that interconnect tumor cells, known as tumor microtubes (TM) that mediate cell-cell communication and probably contribute to resistance to treatment with radiotherapy, chemotherapy, and surgery (Portela et al. 2019). TMs are actin-based filopodia that infiltrate the brain and reach long distances within the brain (Osswald et al. 2015). These TM are associated with a worse prognosis in human gliomas and contribute to the invasion and proliferation, causing effective colonization of the brain by GB cells.

### Genetic basis

The most common genetic lesions in GB patients include mutations in the epidermal growth factor receptor (EGFR), loss of PTEN (PI3K antagonist) and mutation of catalytic subunit of PI3K (Furnari et al. 2007; Louis et al. 2016; Wirsching et al. 2016). Furthermore, it is common to find constitutively active Akt, an important effector of PI3K.

In *Drosophila,* the combination of EGFR and PI3K constitutively active mutations causes a glioma-like condition that reproduces the features of human gliomas, including glial expansion, invasion of the brain, neuronal dysfunction, loss of synapses and neurodegeneration. This model involves the co overexpression of constitutively active forms of *EGFR* (*dEGFR*) and an activated form of the *PI3K* catalytic subunit *p110/PI3K92*k*E* (dPI3K92ECAAX) under the control of *UAS/GAL4*, driven specifically in glial cells via *repo-Gal4.* Coactivation of these signaling pathways in *Drosophila* glial cells increases the levels of MYC, necessary for the tumor transformation and characterized by an increase in the number of glial cells, in the volume of the membrane, reduction in the number of synapses and reduced survival. This coactivation regulates processes such as progression and entry into the cell cycle and protein synthesis (Brand & Perrimon, 1993; Read et al., 2009). So, in this model, MYC plays a central role as it is the point where both routes converge and is essential for tumor transformation.

### Tumor schedule

Among the signaling pathways involved in the progression of GB, the canonical WNT pathway stands out, which is activated after the ligand “Wingless-related integration site” (WNT) binds to the family of Low-density lipoprotein (LRP) or Frizzled (FZD) receptors on the plasma membrane, promoting the expression of cell proliferation genes such as *cyclin D1* and *Myc* (He et al. 1998; Shtutman et al. 1999). The WNT pathway plays a central role in brain development (Loh et al. 2016), adult neuronal physiology (Oliva et al. 2013) and synaptogenesis (Inestrosa & Varela-Nallar 2014; Packard et al. 2002). Alterations in this pathway are associated with neural deficits, in Alzheimeŕs disease (Inestrosa & Varela-Nallar 2014; Marzo et al. 2016) and, above all, GB (Arnés & Casas Tinted 2017; Zuccarini et al. 2018).

Another signaling pathway that has been associated with GB proliferation is the cJun-N-terminal Kinase pathway (JNK) and is currently a drug target for GB (Matsuda et al. 2012). The JNK pathway includes a mitogen-activated protein kinase (MAPK), which belongs to the group of protein kinases stress-activated kinases (SAPKs), a group of kinases that can be activated by any stimulus internal or external that causes cellular stress. The MAPK cascade triggers dual phosphorylation of cytosolic JNK and initiates the phosphorylation of cytoplasmic and nuclear proteins (Chang & Karin 2001), including cytoskeletal and mitochondrial proteins, nuclear transcription factors, protein membrane or nuclear hormone receptors (Bogoyevitch & Kobe 2006). It presents a high homology from *Drosophila* to mammals (Mark & Richardson 2020). In mammals the pathway involves four kinases and mitogens or cytokines that induce MAP3K activation. In *Drosophila,* JNK signaling is initiated by the interaction of the Eiger ligand (Egr), the only member of the TNF ligand superfamily (Igaki et al. 2002; Moreno et al. 2002), with the receptors TNF (TNFR) Grindewal (Grnd), or Wengen (Wng) (Igaki et al. 2009). The ligand-receptor interaction initiates a cascade of phosphorylations (Mark & Richardson 2020). A dual role for the JNK pathway in cell death and survival has been proposed, depending of cell type and context (Mark & Richardson 2020). This double role is especially relevant for CNS pathologies where signals associated with cellular stress increase, such as neurodegeneration and tumorigenesis (Musi et al. 2020; Portela et al. 2019).

In addition, the JNK pathway is the main regulator of *matrix metalloproteases (MMPs)* expression and cell motility in several organisms and tissues, including tumors such as GB (Portela et al., 2019). MMPs are a family of endopeptidases capable of degrading the extracellular matrix (ECM). The members of the MMP family include the “classical” MMPs, the membrane bound MMPs (MT MMP), ADAMs (a disintegrin and metalloproteinase; adamlysins), and ADAMTS (a disintegrin and metalloproteinase with thrombospondin motif). In humans, there are more than 20 members in the MMP family and ADAMTS, including collagenases, gelatinases, stromelysins, some elastases, and aggrecanases (Malemud 2006). There are two orthologues of human MMPs in *Drosophila,* MMP1 and MMP2. MMP1 and MMP2 are anchored to the membrane (Page-McCaw et al. 2003). MMPs are upregulated in various tumors, including GB. Cancer cells produce MMPs that facilitate tumor progression and invasiveness, and upregulation of MMPs in GB is associated with the diffuse growth and has been proposed to play a role in cell migration and infiltration in GB (Nakada et al., 2003). Specifically, among the 23 MMPs present in humans, MMP9, MMP2, and MMP14 are directly involved in the growth and invasion of GB cells (Munaut et al. 2003).

Recent findings in *Drosophila* show that there are three aspects necessary for progression of GB at the cellular level: TM expansion, increased number of GB cells, and loss of synapses in surrounding neurons. GB aggressiveness correlates with progressive activation of the JNK pathway (Portela et al. 2020).

Thus, a timeline has been proposed for the events that occur in the progression of GB in *Drosophila.* Tumor induces infiltration through ultra-long glial membrane protrusions known as TMs, followed by neurodegeneration and GB cell proliferation (Casas-Tintó & Portela 2019; Osswald et al. 2015; Portela et al. 2020). The glial network mediates cell-to-cell communication and promotes the exchange of molecules between cells neurons and GB cells. Consequently, the JNK pathway is activated in GB cells and promotes GB progression and infiltration. Thus, GB cells project TMs that cross the extracellular matrix (ECM) and infiltrate the brain to reach territories distant from the GB primary site (Osswald et al. 2015). In this process, GB cells activate the JNK pathway and accumulate MMPs (MMP1 and 2). These MMPs contribute to the degradation of the extracellular matrix and the expansion of TMs in the brain, these cellular processes facilitate Wg/WNT signaling in GB cells mediated through Frizzled1 receptor (Fz1). Furthermore, Wg/WNT signaling mediates JNK activation in GB cells to continue with the production of MMP and the infiltration process of TM (Portela et al. 2019).

Ultimately, JNK activation in GB cells leads to gene expression upregulation, including MMPs. MMPs mediate ECM degradation facilitating infiltration of TM, communication between cells, and maintaining a positive feedback loop (Portela et al. 2019, 2020). Consequently, MMPs accumulation in GB is an indicator of poor prognosis and the study of MMP-mediated mechanisms is relevant to GB biology and cancer in general (Portela et al. 2020).

GB cells show a progressive increase in JNK pathway activation. JNK pathway is activated in GB cells through the specific receptor Grnd upon binding of Erg ligand from neurons (Portela et al. 2020). This re-localization of Egr to GB cells coincides with the loss of synapses, as an event prior to tumor infiltration and proliferation. Previous studies in *Drosophila* have shown that Egr is expressed in non-tumor brain tissue but accumulates in tumoral cells and activates the JNK pathway. Consequently, GB cells produce MMPs that facilitate infiltration of TM and GB progression (Jarabo et al. 2021; Portela et al. 2019, 2020).

Erg or Grnd deletion experiments in *Drosophila* show that JNK inhibition rescues tumor proliferation and invasiveness (Portela et al. 2020). *In vitro* and *in vivo* experiments showed that JNK inhibitors SP600125 and AS602801 affected GB self-renewal and potential for tumor initiation (Matsuda et al. 2012; Okada et al. 2016), although further studies are required to understand the contribution of JNK pathway and use it as an anticancer target.

In recent years the central role of TM in GB biology has emerged as a fundamental mechanism for the progression of GB, becoming an attractive field of study for possible GB treatments. Furthermore, the JNK pathway is a drug target for GB (Matsuda et al., 2012).

### Drosophila melanogaster as a disease model

DIPG study has mainly used *in vitro* cell models and tumor xenografts in the brain of mice (Lapin et al. 2017; Wu et al. 2014). Although both models are useful for understanding cancer and the search for treatments, have limitations making it necessary to generate a robust model that allows to deepen the investigation of the DIPG.

The use of *Drosophila melanogaster* to reproduce a wide variety of human diseases is becoming more common. *Drosophila* is one of the most effective models to advance in the understanding cancer (Chen & Read 2019). Its short generation time, low costs of maintenance, and the varied availability of genetic tools, allow their use as model for the study of relevant pathways in biomedical research, including cancer (Mirzoyan et al. 2019). In addition, 60% of *Drosophila* genes are conserved in humans, and around 75% of the genes associated with human pathologies, have a functional orthologue in *Drosophila* (Chen & Read 2019)*. Drosophila* glial cells display cellular properties, and physiological functions like human glia. In consequence, *Drosophila* as a disease model presents great advantages crucial for the study of glioma biology (Chen & Read 2019).

On the other hand, *Drosophila melanogaster* has already been validated as a model of human diseases, including GB. The genes most frequently affected in DIPG are conserved in *Drosophila* (Ahmad & Henikoff 2021; Casas-Tintó & Portela 2019; Furnari et al. 2007). All this makes *Drosophila* a suitable experimental model to advance in the understanding of DIPG.

In this report we describe the generation of a new DIPG genetic model in *Drosophila* that recapitulates features of human disease. We have validated this model and used it to determine the molecular mechanisms involved in DIPG progression. In particular, we analyze the expression and contribution of JNK pathway, MMPs and synaptic genes to DIPG progression.

## MATERIALS AND METHODS

### UAS/GAL4 expression binary system

We used the binary expression system UAS/GAL4 in *Drosophila melanogaster* to induce gene expression or gene knockdown. It is based on the GAL4 transcriptional activator from *Saccharomyces cerevisiae* that allows directing its expression to a specific tissue using specific promoters, and the activating sequence UAS *(Upstream Activating Sequence)* that can be fused with a gene of interest. By crossing two parental lines, one carrying the gene of interest fused to the UAS sequence, with another parental line containing the gene encoding GAL4 under the control of a tissue-specific promoter, the offspring expresses the GAL4 activator that upon binding to the UAS sequences, activates the expression of the gene o genes of interest in a specific tissue (Brand & Perrimon, 1993).

In addition, we incorporated a thermosensitive form of the repressor protein Gal80 (Gal80^TS^). It provides temporal control of the expression upon temperature shift. Gal80^TS^ is active at 17°C preventing the binding of GAL4 to the UAS sequence. While, at a temperature of 29°C the GAL80 protein is inactive, so GAL4 binds to the UAS sequence and promotes the expression of the gene of interest (SE McGuire et al., 2003).

The *repo (reverse polarity protein)* promoter sequence was used to direct the expression of *Gal4* to *Drosophila* glial cells (Xiong et al. 1994).

### Drosophila stocks

*w; UAS-dMyc/MKRS (*Eduardo Moreno, Champalimaud Foundtion, Lisbon, Portugal)

*w; UAS-H3K27/Cyo; MKRS/TM6B (*Kami Ahmad, Fred Hutchinson Cancer research Center, Seattle, EEUU)

*w; tub-Gal80ts/Cyo; repoGal4:UAS-myrRFP/MKRS (*BL7108 (tub-Gal80ts); BL 7415 (repoGal4); BL7119 (UAS-myrRFP))

*w; UAS-LacZ (* BL 3955)

*w; ImpL2-MIMIC* (BL-59246)

*w; UAS-ihog-RFP/MKRS* Isabel Guerrero. CBMSO, Madrid)

*w; UAS-Liprin* α*-GFP:UAS D3brp-Strawberry* (Alberto Ferrús, Instituto Cajal, Madrid)

*w; UAS-TRE-RFP/Cyo* (J.P Vincent, Crick Institute, London, UK)

*w; UAS-grndMINOS/Cyo* (P. Leopold, Institut Curie, Paris, France)

### Larval brain dissection

To dissect the larvae, we added phosphate buffered saline (PBS) to a dissection dish. Then we hold the larva with dissecting forceps and cut the underside of the larva. After that, the head of the larva is held with tweezers and, in turn, the other forceps are inserted through the opening of the section. Next, with the tweezers that hold the head, this is pushed until the internal wall and viscera are exposed to the exterior. Finally, the viscera of the larva are removed, the brain is located and separated from the rest of the body.

### Adult brain dissection

First, the flies were anesthetized with CO2 and immobilized on a dissection plate with tungsten pins. A drop of 1X PBS is added and proceed to the dissection: first we remove the proboscis and the cuticle from the head. Once the brain is exposed, the tracheal system is cleaned, and the remains of optical pigment are removed. Finally, the brain is separated from the rest of the body.

### Immunostaining

Fixation with 4% formaldehyde (FA) at room temperature for 20 minutes.

Two consecutive washes of 10 minutes and one wash of 30 minutes with PBT (PBS 1X + Triton at 0.3%) under stirring at room temperature. Incubation in blocking solution (0.3% PBT + 5% BSA) under agitation for 30 minutes at room temperature. Incubation with the primary antibody in blocking solution at 4°C overnight. Two consecutive washes of 10 minutes and one wash of 30 minutes with 0.3% PBT under agitation at room temperature. Incubation with secondary antibody (fluorochrome-conjugated) in blocking solution for 2 hours under agitation and in the dark at room temperature. Three consecutive washes of 10 minutes with 0.3% PBT under agitation at room temperature. Mounting of samples on microscope slides, using mounting medium (Vectashield with DAPI), covering the slide with a coverslip and storing at 4°C until the analysis.

### Antibodies (primary and secondary)

| Primary antibody | Origin | Concentration |
| --- | --- | --- |
| Anti-Repo (Mouse) | DSHB (Developmental Studies Hybridoma Bank) | 1:200 |
| Anti-Myc (Guineapig) | FA Martín (Instituto Cajal, CSIC, Madrid, Spain) | 1:100 |
| Anti-MMP1 (Mouse) | DSHB (Developmental Studies Hybridoma Bank) | 1:50 |
| Anti-MMP2 (Mouse) | K. Broadie (Vanderbilt University, Nashville, TN, USA) | 1:50 |
| Anti-HRP (Rabbit) | Jackson ImmunoResearch 111-035-144 | 1:200 |

The anti-Repo antibody recognizes the transcription factor encoded by the *repo* gene, which is used to mark the nuclei of glial cells. The anti-Myc antibody labels the protein dMyc, so it will label the nuclei of those cells in which *dMyc* is expressed. The anti-MMP1 antibody recognizes matrix metalloprotease 1. The anti-MMP2 antibody recognizes matrix metalloprotease 2. The anti-HRP antibody labels the membrane of the neurons in *Drosophila*.

Secondary antibodies (Thermofisher): anti-mouse or anti-rabbit Alexa 488, 568 or 647. DNA was stained with 2-(4-amidinophenyl)-1H-indole-6-carboxami-Dine (DAPI) at 1 μM in Vectashield mounting media (Vector Laboratories).

### Image acquisition and analysis

We used a Leica SP5 confocal microscope (Leica Microsystems) to acquire the images. The images were taken in a 20X and 63X immersion objectives, each 1.5μm in the Z axis. The resolution used was 1024 x 1024 pixels. The lasers used were UV 405 nm, Argon 488 nm, DPSS 561 nm and HeNe 633 nm.

For image analysis, we used IMARIS software (www.bitplane.com) and ImageJ 1.5.1.

The IMARIS program allows to quantify the amount of fluorescent signal from a 3D image taken using confocal microscopy, and to quantify the number of cells that contain a marker, or the volume occupied by the fluorescent signal of said image.

The Image J software allows you to quantify the pixel intensity of the fluorescent signal in an image 3D taken by confocal microscopy and to quantify the amount of MMP present in each region. We analyzed 6 different brains of each genotype. We calculated the average of the pixel intensity values and the dispersion curves.

### Survival and viability assay

We did survival assay with adult males selected from the four genotypes of interest (Control; H3K27MYC; H3K27M and MYC). We used 30 males of each genotype divided into 3 tubes of 10 individuals each. The vials were kept at 29°C in 12h/12h light-dark cycles and 70% humidity. The count of individuals, as well as the passage of surviving flies to new tubes, was done every other day until all subjects died.

For the viability test, the genotypes Control *(UAS-LacZ),* grndMINOS *(UAS-LacZ;UAS-grndMINOS),* H3K27MYC *(UAS-H3K27;UAS-dMYC)* and H3K27MYCgrndMINOS *(UAS-H3K27/ UASgrndMINOS; UAS-dMYC)* all of them carriers of the *repoGal4:UAS-ihog-RFP* construct. The crosses were kept at 25°C, and after 4 days we transferred parental flies to a new vial as a replica. We counted the number of individuals that reached the adult stage every 2 days until no more subjects emerged.

### Statistical analysis

We used GraphPad Prism program for the statistical analysis. We used t-test statistical analysis for the samples in pairs with a normal distribution and the Mann-Whitney test for samples with a non-normal distribution. In the cases in which more than two genotypes were analyzed, we used ANOVA statistical analysis followed by multiple comparison Bonferroni test for samples with a normal distribution; and Kruskal-Wallis test together with the multiple comparison Dunn test for samples with non-normal distribution. For the survival test, we used the Mantel-Cox test.

In all cases, the minimum value chosen to consider a difference as statistically significant was p<0.05.

## RESULTS

### Co-expression of *H3K27* and *MYC* in glial cells increases the number of glial cells and the glial membrane volume in larval stage

To model DIPG we first aimed to determine if the combined expression of *H3K27+MYC* in glial cells reproduced DIPG features in *Drosophila* brains. To validate this model, we used strains of *Drosophila* carrying *UAS-H3K27M and UAS-dMYC* and the transcriptional activator Gal4 expressed in glial cells under the control of *repo* promoter (*repoGal4*). Moreover, we also included the expression of a UAS-construct to mark glial cells with a red fluorescent protein that accumulates in the cell membranes (*UAS-ihogRFP* system). Ihog *(interference hedgehog)* is a membrane protein that mediates the response to Hedgehog signal, it accumulates in cytonemes and on the cell surface (Portela & Casas-Tintó 2020).

As a control, we used strains of *Drosophila repoGal4:UAS-ihog-RFP* crossed with *UAS-LacZ.* This genotype express RFP that accumulates in the glial membrane but does not produce DIPG and was compared with the experimental genotype *H3K27+MYC* (DIPG).

We analyzed *Drosophila* larval brains by immunofluorescence using ant-repo and anti-Myc antibodies which allow the visualization of the glial cells nuclei (repo) and the protein MYC. In addition, the glial membrane volume was visualized by the fluorescent signal of RFP (Figure 1A-B”).

**Figure 1.**
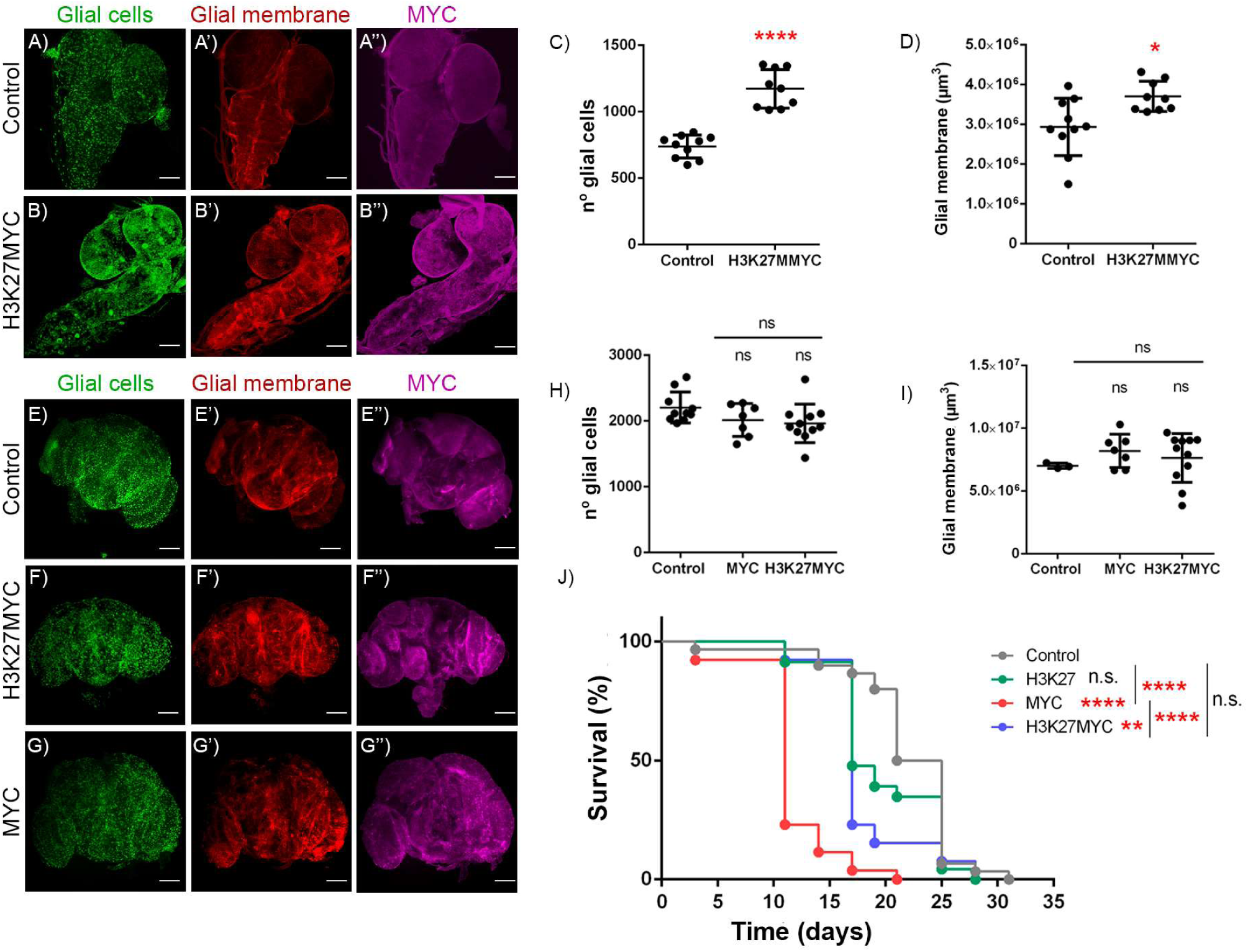
Glial cell co-expression of H*3K27MYC* reproduce the pediatric characteristics of the DIPG. Larvae (A-B) and adult brain images (E-F). Shown in green glia cells nuclei (A, B, E, F, G), in red the glial membrane (A’, B’, E’, F’, G’) in magenta MYC (A’’, B’’, E’, F’’, G’’) in genotypes control (A-A’’, E-E’’), H3K27MYC (B-B’’, F-F’’) and MYC (G-G’). Graphic representation of the number of glial cells (C y G) and glial membrane volume (D y H) in larvae (C y D) and adult (G y H) brain of the genotypes control (n=10 y n=9, respectively), H3K27MYC (n=9 y n=10, respectively) y MYC (n=7). Graphic representation of the survival percentage in adult individuals (J) in genotypes control (grey), H3K27 (green), MYC (red) y H3K27MYC (blue) (n=30). Scale 100μm. * p < 0,05; ** p < 0,001; **** p < 0,00001.

The statistical analysis of the results obtained by 3D reconstruction of the confocal microscopy images showed significant differences between control and DIPG samples in the number of glial cells and in the total volume of glial membrane (Figure 1C, D). These results indicate that the expression in glial cells of *H3K27+MYC* combination increases the number of glial cells and the volume of the glial membrane in the larval stages of *Drosophila*.

### Co-expression of *H3K27and MYC* in glial cells during adult stages does not affect the number of glial cells or glial membrane volume

DIPG is classified as a pediatric-type glioma, with the highest incidence in the middle childhood, the validation of the experimental model requires that it discriminates the development of the tumor based on the age of the individual, and therefore, that reproduces the characteristics of this pediatric glioma in *Drosophila*.

To generate DIPG cells in adult stages, we took advantage of the Gal80^TS^ repression system (see Materials and Methods). The crosses and selected offspring were kept at 17°C (system UAS/GAL4 inactive) until adult stages. Then, once the flies reach the adult mature stages, we maintained the flies of the progeny 7 days at 29°C (active UAS/GAL4 system).

We used *UAS-LacZ* carrying flies together with the *tub-Gal80ts system; repoGal4:UAS-myrRFP* as the control genotype; and flies carrying the combination *H3K27+MYC* combined with the construct *tub-Gal80ts; repoGal4:UAS-myrRFP* as experimental genotype. The myristoylated form of RFP (myrRFP) accumulates in the cell membrane and is expressed in the glia under the control of *repo-Gal4*.

The analysis of adult brains by immunofluorescence using specific antibodies, allowed the visualization of glial cells and MYC expression (anti-repo and anti-MYC), in addition we visualized the membrane of glial cells by myrRFP fluorescence (Figure 1E-G”).

The quantification and statistical analysis of the results did not show significant differences regarding the number of glial cells nor glial membrane volume between both genotypes in adult brains (Figure 1H, I). Therefore, we did not detect significant differences upon the co-expression of *H3K27+MYC* in adult glial cells in the number of glial cells nor to the volume of the glial membrane.

### Co-expression of *H3K27+MYC* in glial cells does not cause premature death

To determine the life span of this genetic *Drosophila* model of DIPG, we analyzed survival and life span. First, we quantified the survival of young larval stages and the transition to adult stages. We counted the number of larvae that reached adult stages upon expression of *H3K27+MYC* or each component alone. The results indicate that *H3K27*+*MYC* expression in glial cells during developmental stages causes a 100% of juvenile death, significantly different compared with *H3K27* or *MYC* expression separately in glial cells. This result indicates that the combination of *H3K27+MYC* expression provokes premature death in early developmental stages, comparable to DIPG in patients.

To further validate the DIPG model, we analyzed the life span of adult individuals. The survival assay was performed in adult individuals of the Control *(LacZ), H3K27, MYC* and *H3K27+MYC* expressed under the control of the *tub-Gal80ts; repoGal4:UAS-myrRFP* system. We maintained the flies at 17°C during all development, and in adult stages, we switched the temperature to 29°C to activate the Gal4/UAS expression system.

The results show that the expression of *MYC*+H3K27 does not reduce the life span of adult flies compare with those expressing *MYC* or *H3K27* alone (Figure 1J). Therefore, the co-expression of *H3K27+MYC* does not synergize and does not cause premature death in adult flies.

### DIPG does not increase JNK pathway signaling

The JNK pathway is upregulated in various tumors, including GB. Recent studies in *Drosophila* show that activation of JNK through Grnd receptor is necessary for GB progression (Portela et al., 2019). In consequence, we analyzed if JNK activation is associated with tumor progression in this model of DIPG.

To determine if the JNK pathway is activated in DIPG cells, we used the previously validated *Tre-RFP* reporter, whose transcription is specifically activated in response to JNK signaling (Portela et al. 2019). The analysis and quantification of *Tre-RFP* positive cells in the brain did not show significant differences between the Control and H3K27+MYC samples (Figure 2A-C). These results suggest that the co-expression of *H3K27* and *MYC* in glial cells does not increase JNK signaling, unlike what occurs in GB cells.

**Figure 2.**
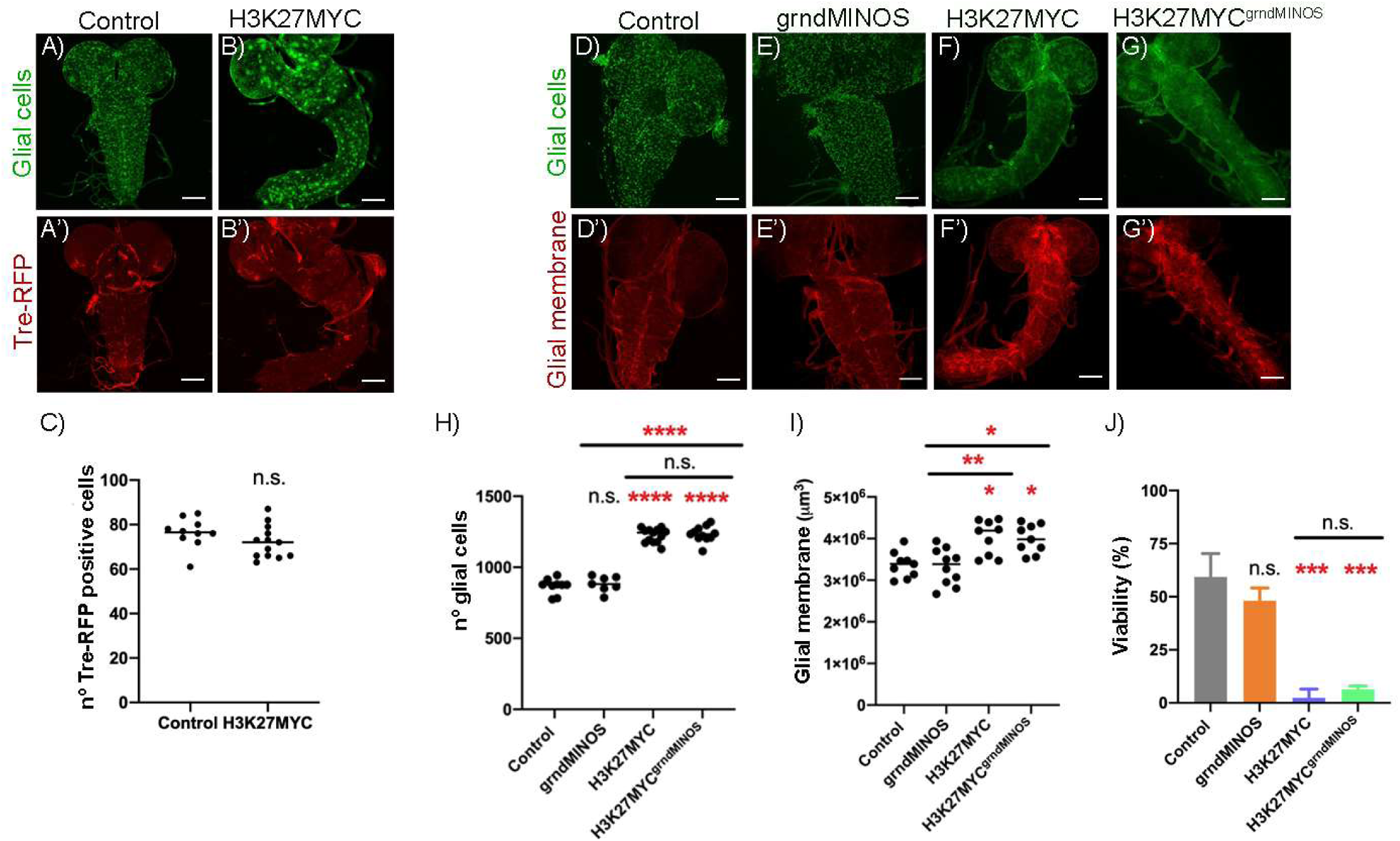
Glial cell co-expression of *H3K27MYC* does not increase JNK signaling. Larvae brain images (A-B, D-G). Shown in green glial cell nuclei (A, B, D, E, F, G), in red the JNK reporter Tre-RFP (A’ y B’) and the glial membrane (D’-G’) from the genotypes cntrol (A-A’, D-D’), H3K27MYC (B-B’, F-F’), grndMINOS (E, E’) y H3K27MYC grndMINOS (G, G’). Graphic representation of the number of Tre-RFP positive cells (C) in control (n=10) y H3K27MYC (n=12), number of glial cells (H) and glial membrane volume (I) in control (n=9), grndMINOS (n=10), H3K27MYC (n=9), H3K27MYC grndMINOS (n=9). Graphic representation of the larvae viability percentage (J) in the genotypes control (grey), grndMINOS (orange), H3K27MYC (blue) y H3K27MYC grndMINOS (green). Scale 100μm. * p < 0,05; ** p < 0,001; *** p < 0,0001; **** p < 0,00001.

*Erg* or *Grnd* deletion experiments in *Drosophila* show that JNK inhibition rescues tumor proliferation and invasiveness in GB models (Portela et al. 2020). To confirm that JNK signaling is not involved in the progression of DIPG, we reduced JNK signaling using flies that express the dominant negative form of Grnd (*UAS-grndMINOS*) (Portela et al. 2020). We analyzed fundamental aspects of the DIPG including the number of glial cells and the volume of the glial membrane (Figure 2D-G). The results did not show significant differences in the number of glial cells or the volume of glial membrane between *H3K27+MYC* and *H3K27+MYC+grndMINOS*. In addition, we observed significant differences in both variables between the control and *H3K27+MYC* genotypes; and control and *H3K27+MYC+grndMINOS* (Figure 2H-I). Therefore, the blockage of JNK signaling does not modify these fundamental aspects of DIPG cells biology, supporting that, unlike in GB, JNK signaling in glial cells is not required in DIPG tumor progression.

Likewise, we did a larval viability assay to further determine the contribution of JNK pathway. We compared the viability of control genotypes, *H3K27+MYC*, *grndMINOS* and *H3K27+MYC+grndMINOS*. The quantification of the percentage of individuals that reached the adult stage showed significant differences between the control and *H3K27+MYC* genotypes, and between control and *H3K27+MYC+grndMINOS*. However, we did not find significant differences between *H3K27+MYC* and *H3K27+MYC+grndMINOS* (Figure 2J). Thus, the inhibition of JNK signaling in glial cells does not rescue the reduction of life span caused by DIPG progression, concluding that JNK signaling is not determinant for the progression of the DIPG.

### DIPG cells increase MMP protein accumulation

A fundamental aspect of DIPG is its ability to infiltrate the brain, a feature in common with GB. Previous studies revealed that GB cells produce MMPs that mediate the degradation of the ECM, thus favoring the infiltration and progression of the tumor (Malemud 2006; Nakada et al. 2003; Portela & Casas-Tintó 2020; Portela et al. 2020).

To determine if MMPs proteins increase in DIPG model cells, we stained larval brain samples for MMP1 and MMP2 of the following genotypes: control (*LacZ*) and *H3K27+MYC* (Figure 3A-F). The analysis of the average pixel intensity of the signal corresponding to MMP1 and MMP2 (Materials and methods) in different regions of the ventral ganglion of larva brain revealed a significant increase in *H3K27+MYC* samples compared to control in both cases (Figure 3C-G). So, DIPG cells increase the amount of MMP1 and MMP2 proteins, although the increase obtained in the case of MMP1 (Figure 3C) was higher than in the case of MMP2 (Figure 3G).

**Figure 3.**
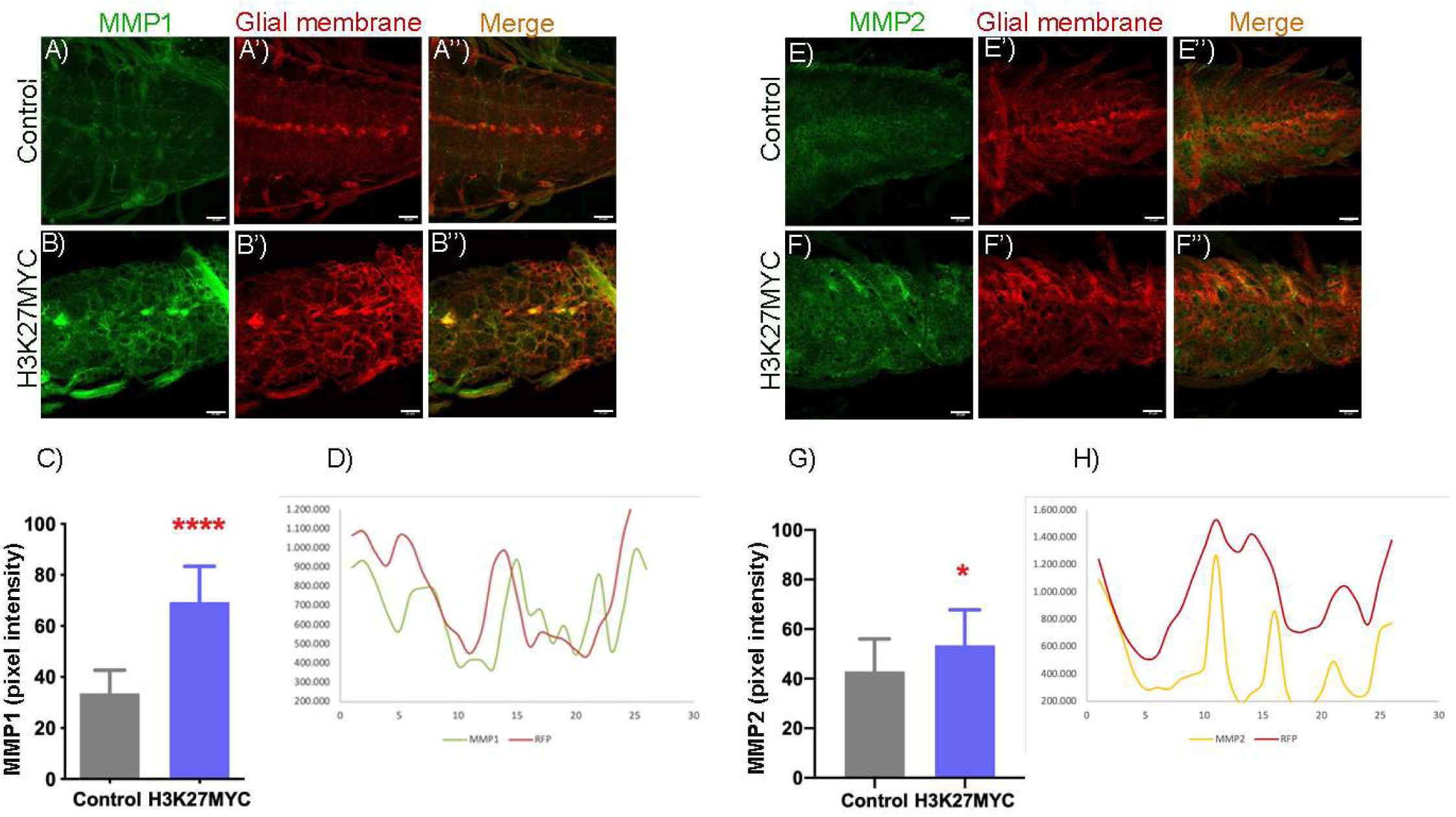
Glial cell co-expression of *H3K27+MYC* increases the levels of MMP1 and MMP2. Larvae brain ventral ganglion images (A-F) of the genotypes control (A-A’’, E-E’’) and H3K27MYC (B-B’’, F-F’’). Shown in green MMP1 (A, B) y MMP2 (E, F), in red the glial membrane (A’-F’) and in orange the fusion of both channels (A’’-F’’). Graphic representation of the amount of MMP1 (C) and MMP2 (G) measured as the mean pixel intensity in the green (MMP1 y MMP2) and red (RFP) channel in 6 different regions of the ventral ganglion of each brain, length 6, in control (n=6) and H3K27MYC (n=7). Graphic representation of the dispersion curves of the pixel intensity values (D-H) of MMP1 (green), MMP2 (yellow) and RFP (red) in the ventral ganglion of larvae brains H3K27MYC. Scale 25μm. * p < 0,05; **** p < 0,00001.

Finally, we determined the relative concentration of MMPs with respect to DIPG cells membrane. To visualize DIPG cells membrane we used the fluorescence of the myrRFP (Figure 3A’-F’). The graphical representation of MMP and RFP intensity in the ventral ganglion of *H3K27+MYC* larval brains showed MMP1 and MMP2 peaks corresponding to a displaced region with respect to the DIPG cell membrane (Figure 3D-H). These results suggest that MMPs are accumulated in the immediate extracellular region of DIPG cells as occurs in GB cells (Conte et al. 2021).

### DIPG cells accumulate synaptic proteins as GB cells

*Drosophila* models of GB have proved the accumulation of synaptic proteins, including Liprin α and bruchpilot (brp). These proteins are crucial for regulating synaptic function and structure, as well as the assembly and maturation of the presynaptic region (REFERENCE). The formation of intratumoral and neuron-GB synapses is required for GB progression and the silencing of synaptic genes in GB cells prevents the expansion of the tumor and expands the lifespan of GB mouse and *Drosophila* models (Losada-Pérez et al. 2022; Venkataramani et al. 2019; Venkatesh et al. 2019). Therefore, the contribution of synaptic genes to GB progression is significant.

To determine if DIPG cells in flies reproduce the accumulation of synaptic proteins, we used the validated fusion proteins Liprin α-GFP and brp-D3-Cherry expressed in glial cells and DIPG cells under the control of the UAS/Gal4 system (Figure 4A-B”). The membrane of glial cells is also marked in red (RFP) and therefore, we quantified the GFP signal that corresponds to Liprin α. The quantification shows that the expression of *H3K27*+*Myc* in glial cells promotes the accumulation of Liprin α.

**Figure 4.**
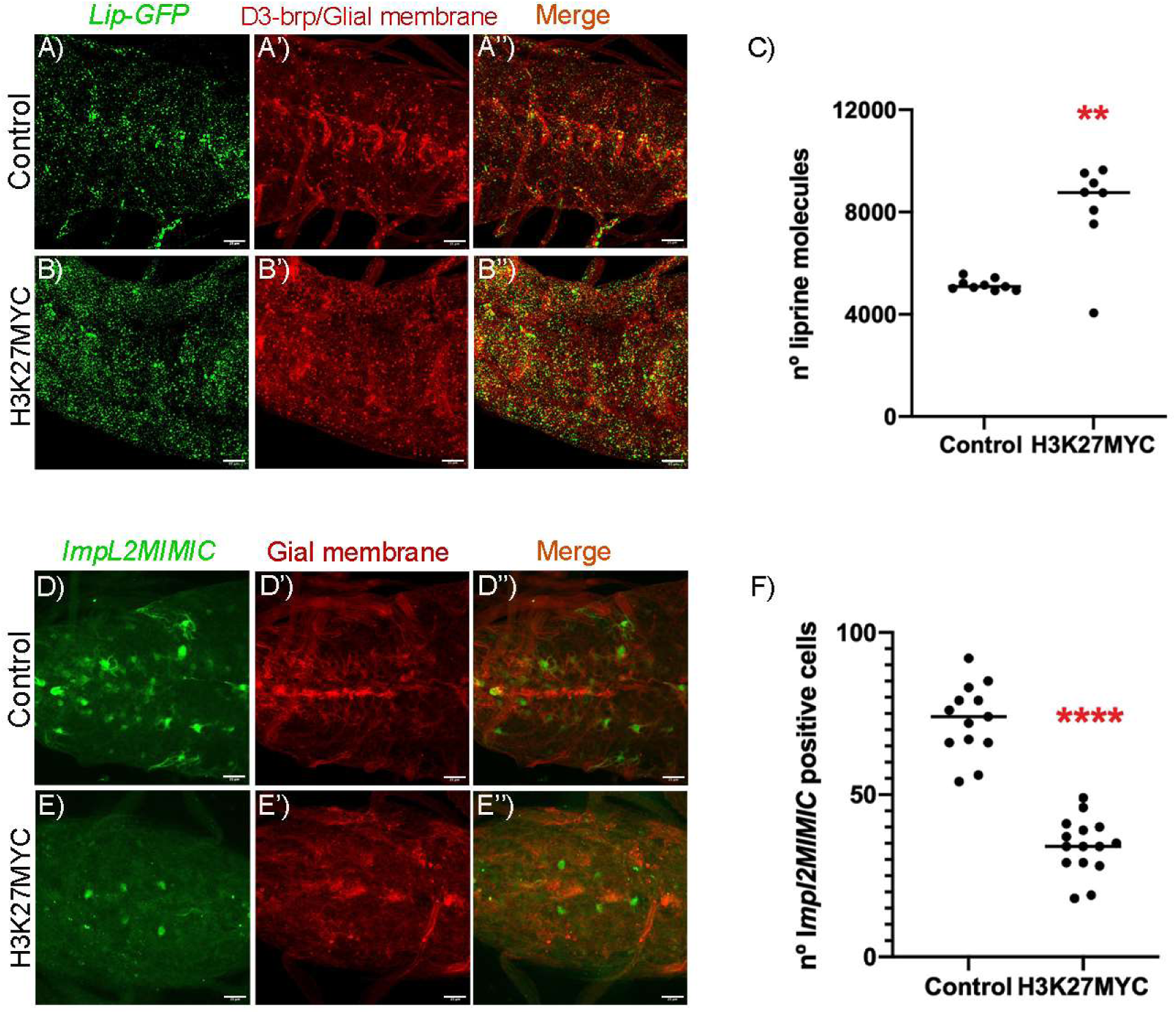
Glial cell co-expressing H3K27MYC accumulate synaptic proteins and do not produce insulin pathway antagonist ImpL2. Larvae brain ventral ganglion images (A-E) of the genotypes control (A-A’’, D-D’’) and H3K27MYC (B-B’’, E-E’’). Shown in green the synaptic protein liprin α (B) marked with the reporter Lip-GFP, and the insulin pathway antagonist ImpL2 (D) marked with the reporter ImpL2-GFP; in red the glial membrane (A’, B’, D’, E’). Graphic representation of the number of liprin molecules in the ventral ganglion (C) in control (n=9) and H3K27MYC (n=8). Graphic representation of the number of ImpL2MIMIC positive cells (F) in control (n=13) and H3K27MYC (n=15). Scale 25μm. ** p < 0,001; **** p < 0,00001.

### DIPG cells do not reproduce GB *ImpL2* expression

ImpL2 is produced by GB cells and mediates the communication from GB cells to neurons. ImpL2 is an antagonist of the insulin pathway produced by GB cells. It attenuates insulin signaling in neighboring neurons which promotes mitochondrial dysfunction and synapse loss in neurons. In consequence, neurons undergo degeneration and GB progress more effectively (Jarabo et al. 2021).

To assess if DIPG cells in *Drosophila* reproduces this behavior, we used a validated MIMIC GFP reporter that replicates *ImpL2* expression (Nagarkar-Jaiswal et al. 2015), in control and *H3K27+Myc* expressing brains (Figure 4D-E”). The quantification of GFP signal in DIPG brains show a reduction that suggest a downregulation of *ImpL2* in DIPG cells. (Figure 4F), opposed to the results described for GB cells.

## DISCUSSION

DIPG is a type of deadly pediatric glioma that currently lacks effective treatments. It has a peak of incidence in middle childhood and a life expectancy of less than 1 year (Lapin et al. 2017; Wu et al. 2014). Therefore, a better understanding of the molecular mechanisms involved in the biology of DIPG is necessary for the development of new therapeutic strategies.

Therefore, the generation of a robust experimental model capable of reproducing the fundamental features of the disease. The most relevant characteristic of this tumor is the presence of mutations in *histone H3*, which is occurs in 80% of cases and is associated with worse outcomes. However, early activation of the H3K27M pathway is not sufficient to produce tumorigenesis, and one of the characteristic features of H3K27M tumors is epigenetic activation of the MYC pathway (Lapin et al. 2017). Therefore, the experimental model used in this work is based on the expression in glial cells of the *H3K27M+MYC* combination.

One of the fundamental aspects of the disease is glial expansion, for this reason we analyzed glial cell number and glial membrane volume in *Drosophila* brain upon expression of *H3K27+MYC* in glial cells. The results show that the co-expression of *H3K27* and *MYC* in glial cells increases cell number and glial membrane volume in the ventral ganglion of larvae. This region is equivalent to the brainstem in humans, where the DIPG originates in patients (Figure 1A-D). Hence, this genetic model reproduces the characteristics of the disease and tumor location. In addition, *Drosophila* DIPG brain samples show abnormally larger glial cells in the ventral ganglion (Figure 1B).

On the other hand, the study in adults reveals that the model reproduces the pediatric characteristic of the tumor. The results obtained reveal that the co-expression of *H3K27+MYC* in the adult brain glial cells does not cause a significant phenotype (Figure 5E-I). Moreover, there are no significant differences in DIPG adult survival compared with *MYC* or *H3K27* expression (Figure 5J). So, H3K27+MYC overexpression flies have a similar life span than *MYC* or H3K27.

Next, we studied the molecular mechanisms involved in the progression and development of the DIPG. Molecular studies showing that DIPG is a unique type of glioma that shares some biological similarities with GB (Jones & Baker 2014). A common feature between both types of gliomas is its infiltrative capacity, recently describe in the study of the role of Tumor Microtubes (TMs). Findings in *Drosophila* show that JNK activation through the Grnd receptor is necessary for GB progression (Portela et al. 2019). Therefore, to determine if the activation of the JNK is implicated in the progression of this DIPG model we used a JNK pathway reporter *(Tre-RFP)*. We analyzed whether the co-expression of *H3K27+MYC* in glial cells increases the signal of Tre-RFP reporter. The quantification of the results showed no significant differences in the number of cells that presented active signaling between control and H3K27+MYC (Figure 2A-C). Thus, the results suggest that there is no activation of JNK in transformed glial cells.

To confirm that the JNK pathway is not involved in the progression of DIPG, we analyzed whether the inhibition of JNK through the reduction of Grnd receptor activity rescues fundamentals aspects of the disease. Analysis of glial cell number and glial membrane volume showed significant differences between the control and *grndMINOS* genotypes, both with respect to *H3K27+MYC*, as with *H3K27+MYC* mutant for Grnd (*H3K27+MYC+grndMINOS*). In addition, we did not observe significant differences between the results obtained in *H3K27+MYC* and *H3K27+MYC+grndMINOS* (Figure 2D-G). Thus, the inhibition of JNK does not rescue the number of glial cells or the volume of the membrane in the model, again indicating that JNK activation does not mediate the progression of tumor. Likewise, the viability test in larvae confirmed these results (Figure 6J). Therefore, the inhibition of JNK nor does it rescue the viability of individuals.

Different studies have shown that MMPs play a fundamental role in the ability to tumor infiltration. These *MMPs* are over-expressed in different types of cancer, including GB where they mediate ECM degradation, favoring tumor progression (Portela et al., 2019, 2020). Given the diffuse nature of the disease, it was evaluated whether, as occurs in GB, there is an increase in the expression of the *MMPs* in DIPG brains that mediate the infiltration of the TM. We visualized through immunohistochemistry the presence of MMP1 and MMP2 in the ventral ganglion of the larval brain. Analysis of pixel intensity revealed-significant differences between the Control and *H3K27+MYC*, both in the case of MMP1 and MMP2, indicating that the co-expression of *H3K27+MYC* in glial cells increases the levels of MMP1 and MMP2 in the ventral ganglion of larval brain (Figure 7A-C, 7E-G). So, just like in GB, MMPs seem to be involved in the progression of DIPG. In addition, the increase obtained in the case of MMP1 is higher than the case of MMP2 (Figure 7C, 7G).

On the other hand, the dispersion curves obtained with the pixel intensity averages of MMP1, MMP2 and RFP in the ventral ganglion of the brain of H3K27MYC larvae, show a displacement of the peak of the MMPs, both MMP1 and MMP2, with respect to that of RFP (Figure 7D, 7H). This suggests that the MMPs are produced by glia and are later secreted into the extracellular space where they mediate ECM degradation. This last hypothesis agrees with the results obtained in the pixel intensity measurements and with studies proposing that MMP1 is secreted while MMP2 as a membrane anchoring sequence (Page-McCaw et al. 2003).

The results obtained suggest that, as in GB, in DIPG the MMPs are involved in the progression of the tumor, probably favoring the infiltration of TMs through the ECM. The study of JNK signaling in this DIPG model indicates that JNK is not required for tumor progression. Which raises a series of questions: what signaling pathway is mediating this increase in MMPs?, could this path present functions analogous to the JNK? Could there be coordination between the two, “exchanging” functions during development?

Besides, our recent publications in GB show that GB cells form intratumoral synapses and also, healthy neurons interact with GB cells through synaptic contacts. Both types of synapses mediate calcium signaling in GB cells and are required for GB progression. DIPG cells in this *Drosophila* models accumulate Liprin α, suggesting that DIPG cells acquire pre-synaptic identity. These results tempt to speculate with the possibility of intratumoral synapses in DIPG and emerge as a potential target to tackle DIPG progression.

Finally, ImpL2 production in GB cells has been described to mediate the decay of surrounding healthy neurons through cell-to-cell communication. This protein binds to the insulin receptor and attenuates the insulin pathway. In consequence, GB cells undergo progressive degeneration. This phenomenon is not reproduced in this DIPG model, suggesting that either *Drosophila* does not reproduce this process, or that neurodegeneration mediated by insulin signaling decay might not be occurring in DIPG cells.

Further studies are required to answer all these questions. A future approach could be to determine if the WNT pathway plays a relevant role in the biology of the DIPG, as MMPs and WNT signaling are part of a positive feedback loop in which the JNK and TMs are involved and promote GB progression. The WNT pathway plays a central role in the development of brain and alterations in this pathway are associated with neural deficits in different diseases, including GB (Arnés & Casas Tintó 2017; Zuccarini et al. 2018). In addition, this route seems to have an essential role in neurological tumors such as medulloblastoma, the most common primary tumor of the CNS in children, and subependymal giant cell astrocytoma (Jozwiak et al. 2007). In various studies describe that WNT induces MMP expression during development and cancer (Lyu & Joo 2005; Page-McCaw et al. 2003) associated with cell migration and metastasis. In fact, the *MMP2* expression in human GB and its infiltrative abilities correlate with Wnt5 (Kamino et al. 2011).

Therefore, the WNT pathway could be responsible for the increase in MMPs levels obtained in this DIPG model. So, unlike what happens in GB, the activation of the WNT pathway could be inducing the expression of MMPs, without the need for JNK activation in DIPG, and therefore WNT would be decisive for the progression of the tumor. To compare DIPG samples of different genetic origins, future studies should compare sequencing studies and the gene profiling of tumor cells, neurodegeneration caused by DIPG progression and possible altered oncogenic pathways.

DIPG is a very complex, lethal, and incurable disease, which has a very limited number of animal models. Therefore, it is necessary to generate a robust model to dig into the knowledge of the DIPG. The results of this work suggest that the co-expression of *H3K27+MYC* in *Drosophila* glial cells is a valid model for the study of DIPG *in vivo,* since it has been proven to reproduce characteristics of the tumor, including glial invasion, tissue specificity and pediatric nature. The study of the underlying molecular mechanisms shows that, as indicated in the literature, DIPG is a unique entity that shares some similarities with GB, but which requires specific therapeutic approaches, for which it is it is necessary to expand the knowledge of the biology of the DIPG.

## Acnowledgements

We thank Dr. Pilar Sánchez Gómez and Dr. Javier García Castro for helpful discussions. We are grateful to Kami Ahmad, the Bloomington Drosophila stock Centre and the Developmental Studies Hybridoma Bank for supplying fly stocks and antibodies, and FlyBase for its wealth of information. We acknowledge the support of the Confocal Microscopy unit at the ISCIII and Dr. Diego Megias for their help with this project. Research has been funded by grant PI22CIII/00062 (Ministerio de Innovacion y Ciencia [MICINN]) and “Unidos Contra el DIPG” foundation.

